# Patient-Derived Tumor Organoid and Fibroblast Assembloid Models for interrogation of the tumor microenvironment in Esophageal Adenocarcinoma

**DOI:** 10.1101/2024.01.02.572565

**Authors:** Benjamin P. Sharpe, Liliya A. Nazlamova, Carmen Tse, David A. Johnston, Rhianna Blyth, Oliver J. Pickering, Ben Grace, Jack Harrington, Rushda Rajak, Matthew Rose-Zerilli, Zoe S. Walters, Tim J. Underwood

**Author notes:** Correspondence: Tim J. Underwood, School of Cancer Sciences, Somers Cancer Research Building, Tremona Road, Southampton SO16 6YD.

## Abstract

The tumor microenvironment (TME) comprises all non-tumor elements of cancer and strongly influences disease progression and phenotype. To understand tumor biology and accurately test new therapeutic strategies, representative models should contain both tumor cells and normal cells of the TME. Here we describe and characterize co-culture tumor-derived organoids and cancer-associated fibroblasts (CAFs), a major component of the TME, in matrix-embedded assembloid models of esophageal adenocarcinoma (EAC). We demonstrate that the assembloid models faithfully recapitulate the differentiation status of EAC and different CAF phenotypes found in the EAC patient TME. We evaluate cell phenotypes by combining tissue clearing techniques with wholemount immunofluorescence and histology, providing a practical framework for characterization of cancer assembloids.

## Introduction

Esophageal cancer is the sixth leading cause of cancer-related deaths worldwide with more than 500,000 new cases reported annually^1^. Esophageal adenocarcinoma (EAC) is the dominant subtype in Western countries and due to its late-stage presentation and therapy resistance, EAC is a deadly cancer with a 5-year survival of less than 15%. Risk factors include gastroesophageal reflux disease, Barrett’s esophagus, obesity, alcohol and tobacco consumption^2^. Surgery represents the primary treatment for early-stage disease, whereas patients with locally advanced disease are treated with perioperative chemotherapy^3^ or chemoradiotherapy prior to surgery^4^. Despite rapid advances in molecularly targeted treatments in other cancers, progress in EAC has been slow with only HER2 directed therapies entering clinical practice^5^. Most recently immunotherapies, in the form of immune checkpoint inhibitors, have entered the field, but durable responses are rare^6^. The intra- and inter-tumoral heterogeneity of the disease presents an obstacle for introducing new therapies, which may partly be attributed to the lack of physical models that reflect the primary disease.

There is accumulating evidence that the tumor microenvironment (TME) which comprise of components including immune cells, fibroblasts, endothelial cells, adipocytes, and extracellular matrix (ECM) contribute to tumorigenesis as well as therapy resistance^7^. Amongst these, cancer-associated fibroblasts (CAFs) make up a large proportion of the TME^8^. CAFs help control cancer progression in addition to influencing cancer cell proliferation, ECM remodeling, metastasis, and therapy resistance through the secretion of growth factors and ECM^9^. Clinically, CAF-positivity in EAC was associated with worse tumor stage and higher rate of metastasis in addition to shorter disease free and overall survival^10^. We have previously shown that markers of myofibroblast CAF differentiation, α-smooth muscle actin (α-SMA) and periostin (POSTN) are associated with poor prognosis in EAC^8^. Targeting myofibroblast differentiation in EAC CAFs can be achieved in preclinical models using PDE5 inhibitors and sensitizes tumor cells to chemotherapy^11^. These studies highlight the importance of the EAC TME in governing tumor behavior, and the TME must be considered when studying drug sensitivity, which current EAC models are lacking^12^. Close-to-patient models that replicate the cancer-stroma interactions to understand the mechanism of resistance would be beneficial for the development of personalized treatment of EAC.

Patient-derived organoids (PDOs) have a wide range of applications in cancer research including drug testing, personalized or precision medicine and cancer immunotherapies. Cancer organoids are self-organizing cancer cells derived from patient tumors that recapitulate the structure, heterogeneity, histology, and genetic signatures of the primary tumor. PDOs have been successfully established in a variety of cancer types including pancreatic^13^, colorectal^14^ as well as EAC^15^. The major limitation of these PDO models is the lack of TME components which contribute to various hallmarks of cancer and response to therapy^16,17^. To recapitulate the cancer-stroma crosstalk, PDOs may be co-cultured with stromal cells including CAFs^18,19^. Studies in PDAC show that the co-culture of organoid models with pancreatic stellate cells (precursors of CAFs) resulted in the presence of a myofibroblastic and inflammatory CAF populations^20,21^, phenotypes which are observed in PDAC patients^22^. A study by Seino et al. revealed CAFs could supply Wnt to support the growth of a Wnt-non-secreting subtype of PDAC PDO^23^. The addition of CAFs also increased the proliferation and resistance of PDAC PDOs to chemotherapies^19^. In colorectal carcinoma (CRC), CAFs were able to maintain the proliferation of CRC PDO co-cultures and restored key survival pathways and cancer-CAF interactions present in the patient tissue^24^. Additionally, CRC PDO-CAF co-cultures show an enhanced resistance to standard of care drugs^24^. These data are evidence of the crosstalk between the organoids and CAFs and highlights the importance of incorporating CAFs when testing drug responses using PDOs. Whilst EAC PDO models have been established^15^, models that incorporate patient derived CAFs are lacking.

In this study we characterized EAC PDOs and devised a co-culture method using basement-membrane matrix and collagen to enable the co-culture of EAC PDOs with patient-derived CAFs to create novel EAC assembloids^25^. We present an optimized method for non-destructive immunofluorescent staining and imaging of assembloids to facilitate study of CAF-PDO interactions in 3D. We show that these assembloids are a viable approach to model cancer-stromal interactions *in vitro*.

## Results

### Generation of EAC Organoid-Fibroblast Assembloids

To study the interactions of tumor epithelium with cancer-associated fibroblasts, we adapted a protocol established by Seino and colleagues to co-culture pancreatic cancer organoids with fibroblasts^26^. Patient-derived organoids (PDOs) were derived from EAC tissue at resection following representative treatment pathways (neoadjuvant chemotherapy followed by esophagectomy). Fibroblasts were derived and expanded from EAC resection material to generate cancer-associated fibroblasts (CAFs) from EAC tissue via explant outgrowth as previously described^26^.

Grown in 3D culture with basement membrane extract and esophageal organoid growth media, EAC PDOs grow with diverse morphologies ranging from dark and dense to pale and cystic tumor buds (**Figure 1A**). Primary EAC CAFs grown in 2D display a classical elongated, spindle-like morphology (**Figure 1B**). The two cell types were co-cultured in a 2:1 ratio of CAFs:PDOs, starting with 2.5×10^4^ organoid cells and 5×10^4^ CAFs per model (**Figure 1C**). Co-cultures were grown in complete DMEM, avoiding the use of expensive esophageal organoid growth media which contains factors that maintain epithelial stem cell niches^27^ and therefore may affect the phenotype of CAFs in culture. Esophageal PDOs do not survive when grown in basement membrane extract in complete DMEM in the absence of fibroblasts (data not shown), suggesting that CAFs provide factors essential for survival and proliferation of tumor cells *in vitro* as they would *in vivo*^23^. After overnight low-attachment culture conditions facilitating cell aggregation, co-cultures were embedded in a mix of 3:1 rat collagen I:BME2 (basement membrane extract), fed with complete DMEM and cultured for 7 further days *in vitro*. Co-cultures are compact and dense on day 1, initially contracting and then developing round bud-like structures on the periphery of cultures on day 3, which continue to develop until endpoint analysis at day 8 (**Figure 1D**). Fibroblasts grow out of the co-culture and by day 8 they also occupy the surrounding gel.

**Figure 1.**
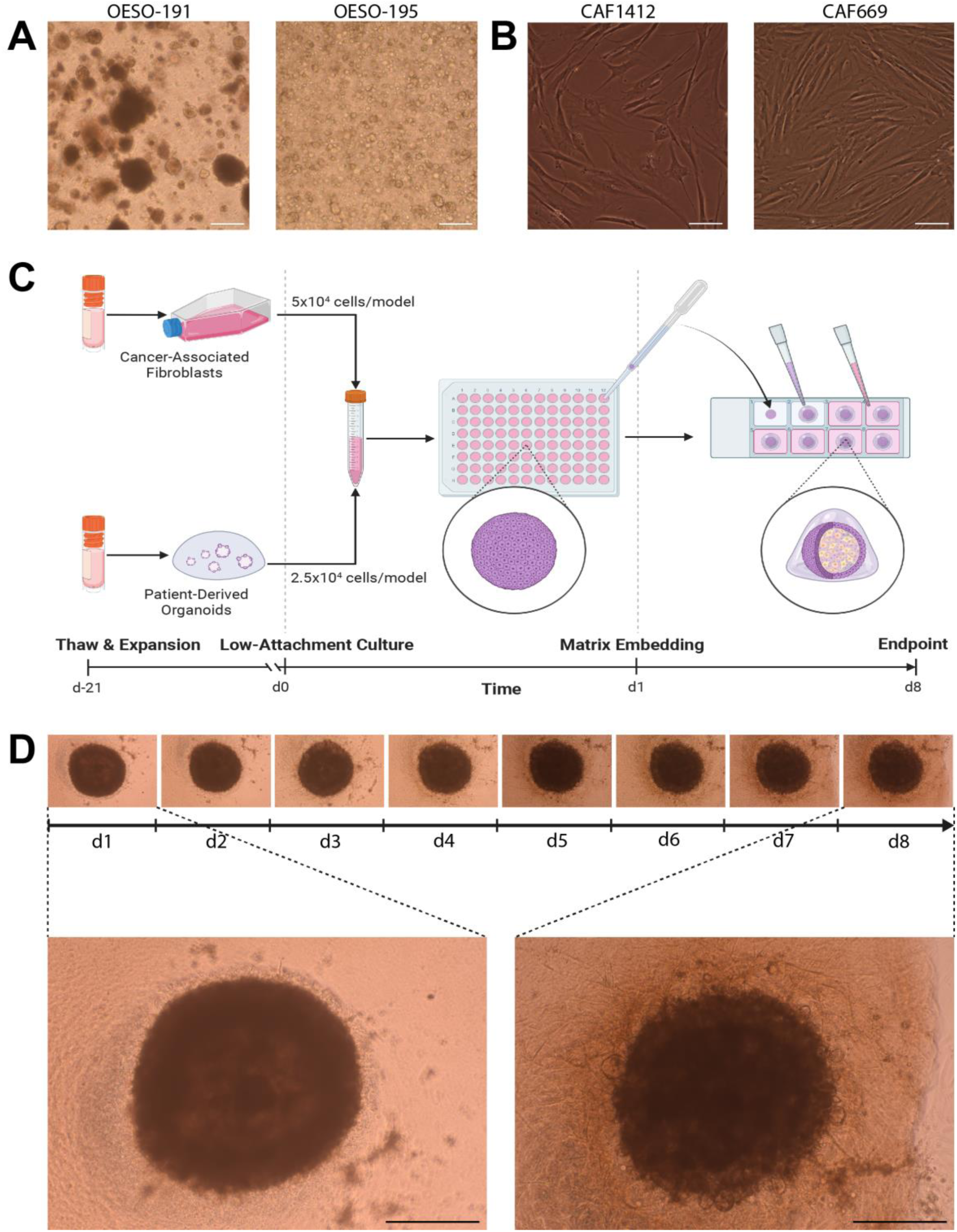
Derivation of primary models and the workflow for creation of EAC assembloids. A. Two example EAC organoids derived from different EAC patients, observed with a 4x objective under phase contrast microscopy. Scale bars – 250µm. B. Two example CAFs derived from different EAC patients, observed with a 20x objective under phase contrast microscopy. Scale bars – 50µm.C. Workflow for production of EAC assembloids. EAC organoids and primary CAFs are expanded prior to model creation, dissociated to single cells, counted and mixed together at a 2:1 ratio of CAFs to organoid cells. The cell suspension is then plated out at 750,000 cells per well in an ultra-low attachment 96-well U-bottom plate. The next day, assembloids are plated in a 3:1 mixture of collagen I:BME2, and complete DMEM media overlayed once set. Assembloids are cultured for a further 7 days prior to harvest. D. Structure formation of EAC assembloids imaged with a 4x objective under phase contrast microscopy every day for 7 days after matrix embedding. Scale bars – 500µm.

### PDO-CAF Assembloids recapitulate features of primary tumors

To verify the epithelial or mesenchymal nature of the input primary tumor cells, we embedded organoids in agarose and subjected them to FFPE tissue processing and immunofluorescence to detect epithelial cytokeratin (using pan-Cytokeratin antibody clone AE1/AE3) and the mesenchymal marker vimentin^28^. Similarly, fibroblasts were seeded onto 8-well chamber slides and stained for pan-cytokeratin and vimentin. EAC organoids were heterogeneously positive for epithelial cytokeratins but negative for vimentin (Figure 2A), and fibroblasts stained conversely (Figure 2B), consistent with their distinct epithelial and stromal origins.

**Figure 2.**
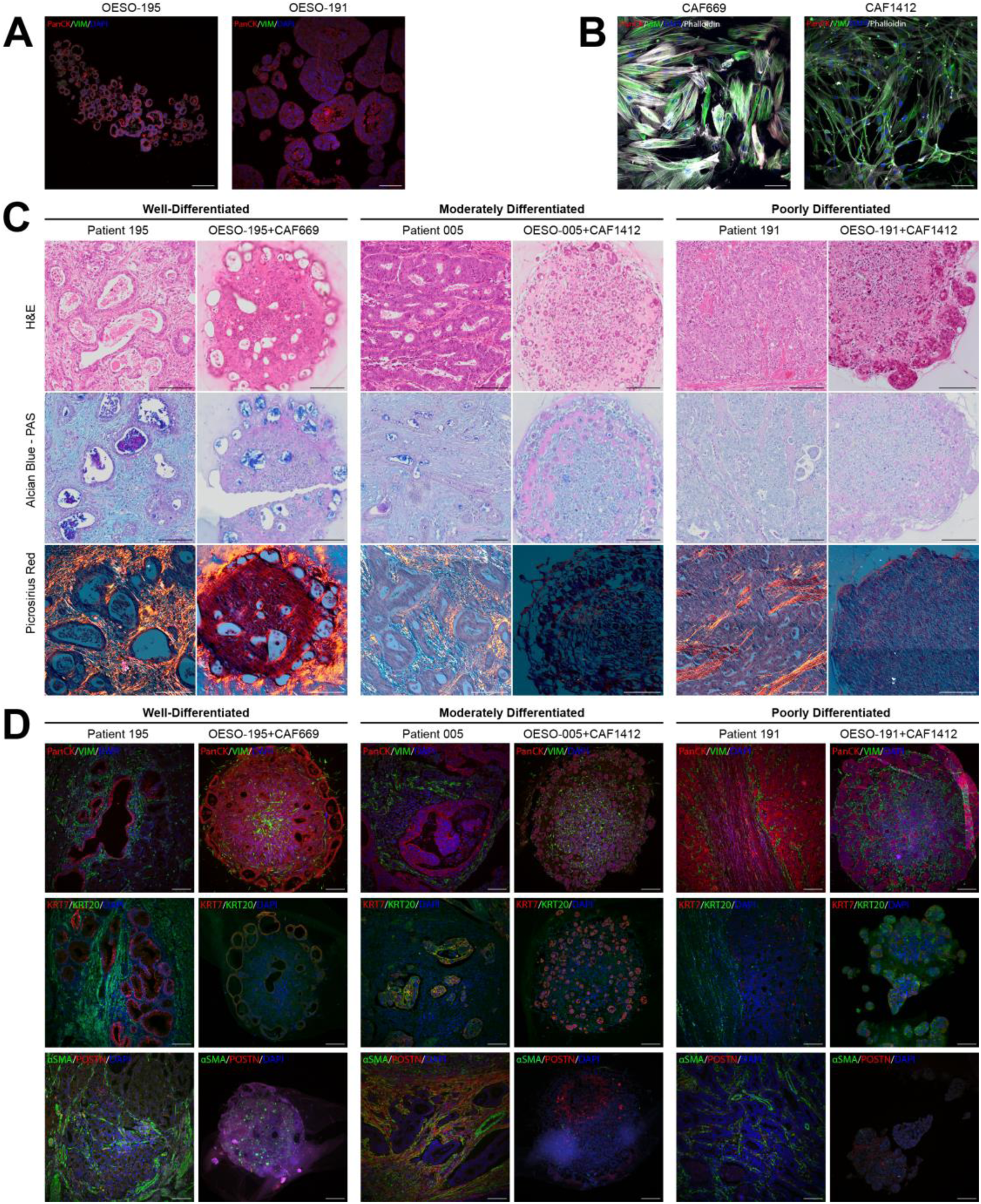
Histological and immunohistochemical characterisation of EAC assembloid phenotypes compared to the corresponding patient tumors. A. Immunohistochemistry of 5um organoid sections grown in monoculture, stained for pan-cytokeratin (red), and vimentin (green). Nuclei were counterstained with DAPI (blue). B. Immunocytochemistry of primary fibroblasts grown in monoculture, stained for pan-cytokeratin (red) and vimentin (green). Actin cytoskeleton is counterstained with phalloidin (grayscale) and nuclei were counterstained with DAPI (blue). Scale bars – 100um. C. 5um sections of EAC assembloids and resected patient tumors stained with tinctorial stains for overall histology (H&E), mucin secretion (Alcian Blue – PAS, mucins in blue/purple), and collagen fibers (Picrosirius Red, collagen in orange) viewed under polarised light. D. Immunofluorescence of EAC assembloids and resected patient tumor sections stained for markers of epithelial and fibroblast cell identity – pan-cytokeratin and vimentin; keratin 7 (glandular epithelium) and keratin 20 (colonic epithelium); and α-SMA and POSTN (myofibroblast differentiation) . Nuclei are counterstained with DAPI. Scale bars – 150um.

We grew three different assembloids containing combinations of three different organoids and two different CAFs for 8 days and processed these for conventional FFPE histology alongside FFPE parental tumor material from esophagectomy. We performed a panel of tinctorial stains to assess the histological composition of the assembloids compared to the parent tumor: H&E staining to assess histological differentiation; alcian blue and periodic acid Schiff (AB-PAS) for the detection of mucins commonly secreted by EAC tumors; and picrosirius red (PSR) to detect collagen fiber deposition and alignment under polarized light (Figure 2C).

Histopathological evaluation of H&E stains revealed similar histological differentiation between the assembloids and parent tumors, with additional differences in histology of assembloids between central and peripheral regions (Figure 2C). Patient 195 had a well-differentiated adenocarcinoma with large hollow tumor acini with intraluminal accumulation of acidic (stained blue) and neutral (stained purple) mucins. Assembloids of organoid OESO-195 with CAFs produced similar phenotypes under H&E and AB-PAS stains, with accumulation of tumor acini on the peripheral regions and accumulation of necro-inflammatory debris in the central region of the assembloid. This is suggestive of central hypoxia and an organization of proliferating cells on the outer layers while cell death occurs centrally. Collagen deposits stained with PSR (orange birefringence) were visible surrounding the outer layers of assembloids containing OESO-195 and CAF669 but were not visible in assembloids containing CAF1412 (Figure 2C). Unlike parent tumors, no assembloids displayed mature collagen networks with aligned collagen fibers, suggesting an absence of collagen-secreting fibroblast phenotypes in these assembloids.

Patient 005 had a well-to-moderately differentiated adenocarcinoma containing dense tumor acini and smaller lumens with acidic mucin accumulation under AB-PAS stain (Figure 2C). Assembloids of organoid OESO-005 displayed moderate differentiation with smaller acinar structures than the parent tumor, and acidic mucin-containing lumens distributed throughout the culture. The central region of the assembloid also contained singly dispersed tumor cells with more poorly differentiated morphology than the peripheral region. Although the corresponding patient tumor forms much larger acini, the overall differentiation status is equivalent to the assembloid, being moderate to well-differentiated.

Patient 191 had a poorly differentiated adenocarcinoma with disorganized tumor nests, notably absent lumens, and no mucin secretion detectable by AB-PAS (Figure 2C). Similarly, assembloid culture of organoid OESO-191 displayed poor differentiation with solid tumor nests in the peripheral region and singly dispersed cells with signet ring cell-like morphology in the central region. Signet ring cell-like morphology indicates intracellular accumulation of mucins which push aside the nucleus, and usually indicates poor clinical behavior. Interestingly, sections derived from the parent tumor did not have observable signet ring histology which may suggest an additional effect of assembloid culture conditions on phenotypic heterogeneity at the histological level.

To ensure that the cancer and fibroblast cell types present in the assembloid were representative of patients, we next performed duplex immunofluorescence on the same FFPE blocks from parent tumor and matched assembloids (Figure 2D). Staining for pan-cytokeratin and vimentin suggested that vimentin-positive fibroblasts surrounded cytokeratin-positive tumor acini in assembloids as they do in the parent tumor. The central region was particularly fibroblast-dense and cytokeratin-positive tumor acini preferentially localized to the periphery, especially in the well-differentiated OESO-195 assembloid. EAC tumor cells are thought to be cytokeratin 7-positive (CK7) but are usually negative for cytokeratin 20 (CK20)^29^. Assembloids mirrored the cytokeratin expression patterns of their parent tumors (Figure 2D). Patient 195 was CK7+/CK20- and the corresponding assembloid reflected this; patient 005 was heterogeneously positive for both CK7 and CK20; whereas the assembloid was mostly CK7+/CK20-with a minority of CK7+/CK20+ tumor cells. Patient 191 had lower levels of CK7 expression and the OESO-191 assembloid contained an admixture of CK7+ and CK20+ tumor cells. Next, we examined the expression of myofibroblast CAF markers α-SMA and POSTN as these are known to be associated with poor prognosis in EAC^8,11^. Assembloids containing OESO-005 and OESO-191 grown with CAF1412 produced an SMA-negative and POSTN-low stromal microenvironment (Figure 2D), whereas the stromal microenvironment of OESO-195 grown with CAF669 produced a stromal microenvironment with SMA-positive cells distributed throughout, suggesting some myofibroblast CAF differentiation in this model.

### Wholemount immunofluorescence demonstrates organization of fibroblasts and morphological variation between patient co-cultures

3D models provide the most information where 3D context and gene/protein expression can be related. However, thick specimens limit diffusion of staining reagents and penetration of light, in part due to presence of lipids^30^. We developed a non-destructive protocol for wholemount immunofluorescence that permits penetration of antibodies and visualization up to ∼250µm depth into the assembloid models (Figure 3A). We adapted this method from Dekkers et al.^31^, using FLASH antigen retrieval from Messal et al.^32^ to improve antibody penetration and imaging depth for large models. The staining protocol takes 5 days and stained specimens can be stored in clearing buffer at 4°C for >2 weeks prior to imaging. Briefly, excess collagen gel was dissected from the assembloids using needles, and we performed antigen retrieval using FLASH reagent 2^32^ for two hours at 37°C. This is an SDS and zwitterionic detergent-based method to remove tissue lipids and improve reagent and light penetration through the specimen. To indirectly immunolabel co-cultures, we incubated with primary antibodies for two days at 4°C, and then for a further two days with secondary antibodies and DAPI for nuclear counterstaining. To clear the assembloids for imaging, we immersed them for at least 30 minutes in fructose-glycerol clearing buffer^31^ (60% (v/v) glycerol and 2.5M fructose, Refractive Index=1.47) and then imaged them with a laser-scanning confocal microscope. To demonstrate the utility of this method, we wholemount stained assembloids with anti-pan-cytokeratin and anti-vimentin to visualize both compartments in native 3D context. Budding tumor structures visible by brightfield observation are revealed in detail when imaged by wholemount IF, showing discrete pan-Cytokeratin-positive structures on the periphery of assembloids (Figure 3B, **Supplementary Movies S1-3**), with internal lumens in the well- and moderately differentiated assembloids and solid tumor masses in poorly differentiated co-cultures. Vimentin-positive fibroblasts can be seen in close association with tumor acini and areas of denser fibroblast presence are more apparent in maximum-intensity projections of wholemount IF stained samples than in paraffin sections. Proliferation of cancer cells within assembloids became apparent when wholemount staining with anti-Ki-67, with poorly differentiated assembloids having the greatest abundance of proliferating cells (Figure 4).

**Figure 3.**
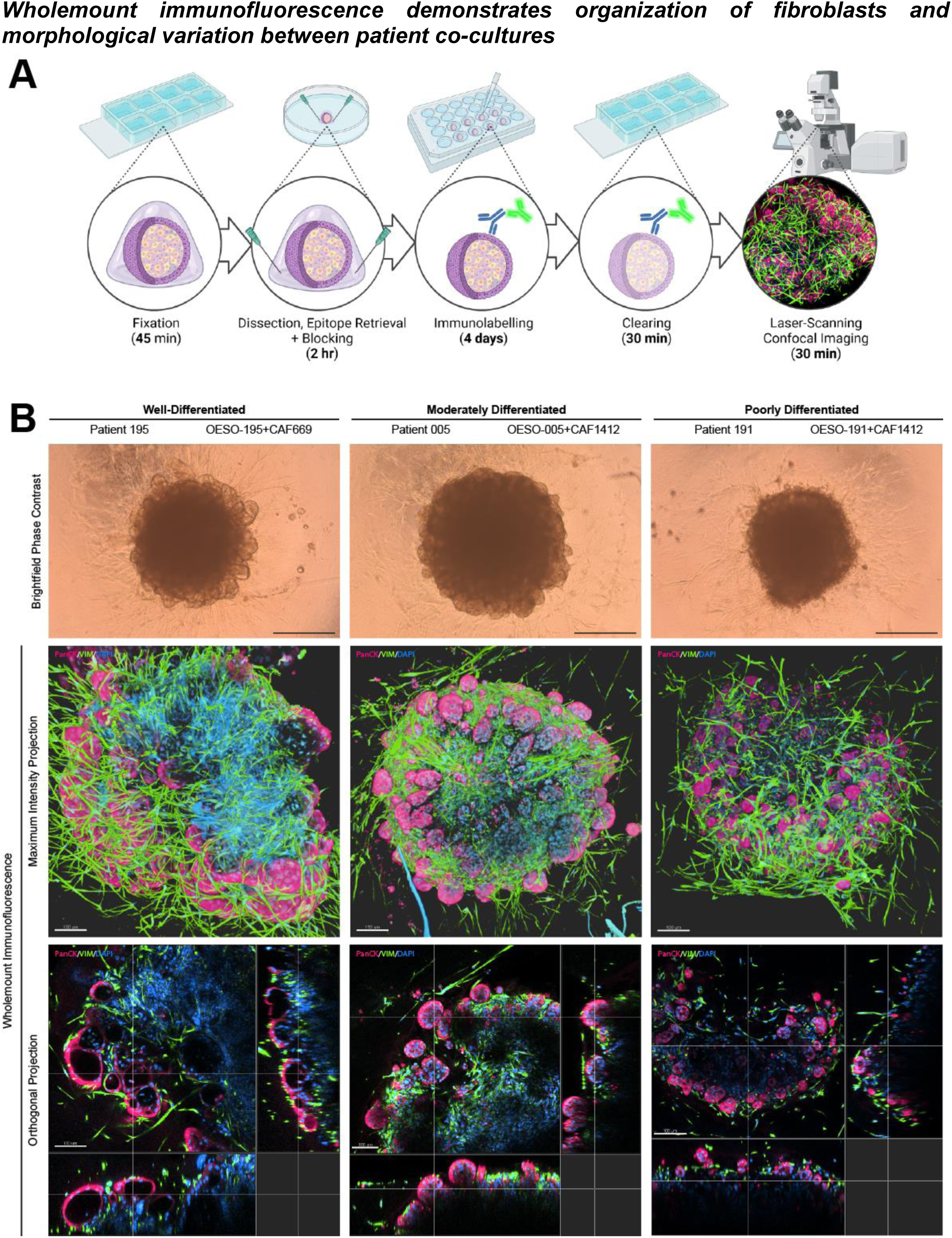
Wholemount immunofluorescent protocol for 3-dimensional visualization of EAC assembloid organization. A. Workflow of the 5-day wholemount staining protocol. EAC assembloids are fixed in 4% PFA, excess collagen is removed to improve antibody penetration, and then assembloids are stained with primary and fluorescent-conjugated secondary antibodies (with DAPI) for 2 days each. Assembloids are cleared in fructose-glycerol clearing buffer and then imaged with a fluorescent laser-scanning confocal microscope. B. Example assembloids from each organoid/CAF combination at day 8 of culture imaged in brightfield phase contrast (scale bars – 500um). Shown below are corresponding Z-projections of wholemount stained and cleared assembloids for pan-cytokeratin (organoid cells, red) and vimentin (fibroblasts, green) in top-down maximum intensity projections and orthogonal projections cutting through outer epithelial buds (scale bars – 100µm unless indicated otherwise). Rendering was performed in Imaris under blend mode. See also Supplementary Movies S1-3.

**Figure 4.**
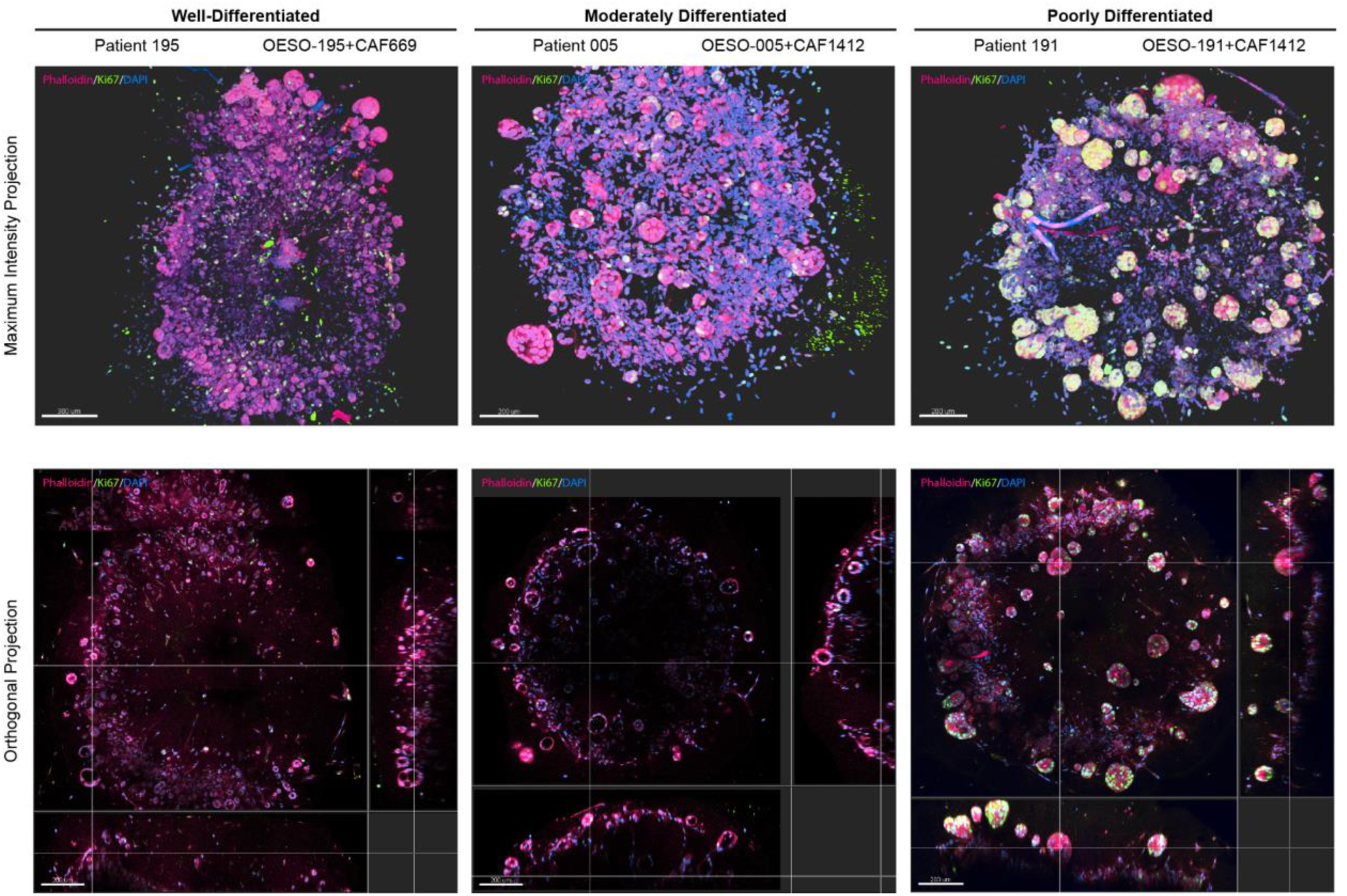
Visualization of cell proliferation in 3D using wholemount immunofluorescence. Z-projections of wholemount stained and cleared 8-day-old assembloids for Ki-67 (proliferating cells, green) counterstained with Phalloidin-iFluor594 (cell bodies, red) in top-down maximum intensity projections and orthogonal projections cutting through outer epithelial buds (scale bars – 200µm). Rendering was performed in Imaris under blend mode.

## Discussion

Organoid cultures preserve the heterogeneity and molecular characteristics of the primary tumor and facilitate testing treatment strategies and understanding responses to them in EAC^15^. However, current models do not incorporate stromal elements of the TME, which provide context cues by both paracrine and juxtacrine interactions. Furthermore, it is well recognized that stromal cells, including CAFs, play a major role in tumor progression and drug response^33^. Assembloids are an emerging technology for the study of epithelial cells within their mesenchymal niches, with assembloid models recently being developed for modelling stromal interactions in the murine normal and malignant colon^34^. Our model combines the versatility of patient-derived tumor organoid lines with well-characterized genomic and transcriptional landscapes, with primary CAFs from heterologous donors. This method facilitates physical contact of CAFs with tumor cells where juxtacrine signalling can take place, as opposed to other methods where fibroblasts loosely associate due to being mixed into the extracellular matrix^35^. The models are low-to-medium throughput, technically simple to produce, require no specialized equipment and would accommodate addition of other cell types to suit the research question (e.g., immune or vascular cells).

We show that EAC assembloids recapitulated features of the primary tumor including histology and differentiation status whilst recapitulating tumor-stroma interactions. We observed that CAFs support the survival and proliferation of EAC PDOs in complete DMEM, crucially enabling low-cost model production and scalability. This was also observed in PDAC^23^, normal colon^34^ and CRC^24^ where growth of the epithelial compartment was supplemented by fibroblasts even in the absence of exogenous Wnt and BMP signaling activators. We observed spatial arrangements of EAC assembloid models comparable to those seen in multicellular tumor spheroids, where reduced access to oxygen and nutrients internally causes models to proliferate in peripheral areas, with central quiescence and necrosis as spheroids grow larger^36^. Progressive growth of tumor buds from the periphery of EAC assembloids and central necro-inflammatory debris suggested a similar dynamic occurring in these models. Interestingly, histology revealed the presence of singly dispersed tumor cells and signet ring cells in central zones of EAC assembloids, which are features associated with poor prognosis in tumors^37^. These did not match with patient tumor histology and, given the large size of the models, we suggest they might be generated by hypoxia within the dense fibroblast-rich core. Hypoxia has previously been reported to push CAFs towards an inflammatory phenotype when co-cultured with mouse PDAC organoids, while leaving myofibroblast markers unaffected^38^. Simulating a hypoxic environment using assembloids offers the opportunity to better understand the impact of hypoxia on fibroblast and organoid phenotypes, and the crosstalk between them.

CAFs are recognized to be heterogenous with different subtypes and different functions in the TME intratumorally^39^.

Our study is limited due to the CAFs being heterologous, therefore not fully mimicking CAF-PDO interactions as they are present in the patient from which the organoid was derived. However, we have shown that CAFs maintain phenotypes observed in monoculture when grown in assembloids. This provides the opportunity to study the contribution of diverse CAF phenotypes from different patient subgroups to tumor cell behavior. Future work aims to use matched EAC PDOs and CAFs to develop patient-specific models as a prediction of drug response which will be correlated to clinic as well as explore the incorporation of matched immune cells to develop a co-culture model for testing immunotherapies. Furthermore, the current method is low throughput, being able to generate <96 models in one experiment and limited by manual handling. To facilitate higher throughput screening efforts which include stromal populations, miniaturization and reducing manual transfer steps in matrix embedding would be required.

In this study we successfully established a disease relevant co-culture model consisting of patient derived CAFs and organoids that enables technically straightforward modelling of tumor-stromal crosstalk. We combined the versatility of patient-derived tumor organoid lines with well-characterized genomic and transcriptional landscapes, with primary cancer-associated fibroblasts from heterologous donors. We developed a non-destructive and inexpensive clearing and imaging toolkit that enables visualization of interactions in co-cultures in 3-dimensions.

## Materials and Methods

### Ethics & Patient Cohort

Patient tumor material was obtained subject to informed consent and local ethical approval (University of Southampton ERGO number 45334) and NHS ethical approval (REC number 18/NE/0234). Key clinical information for each patient involved in organoid and fibroblast derivation is listed below (**Table 1**).

**Table 1.**
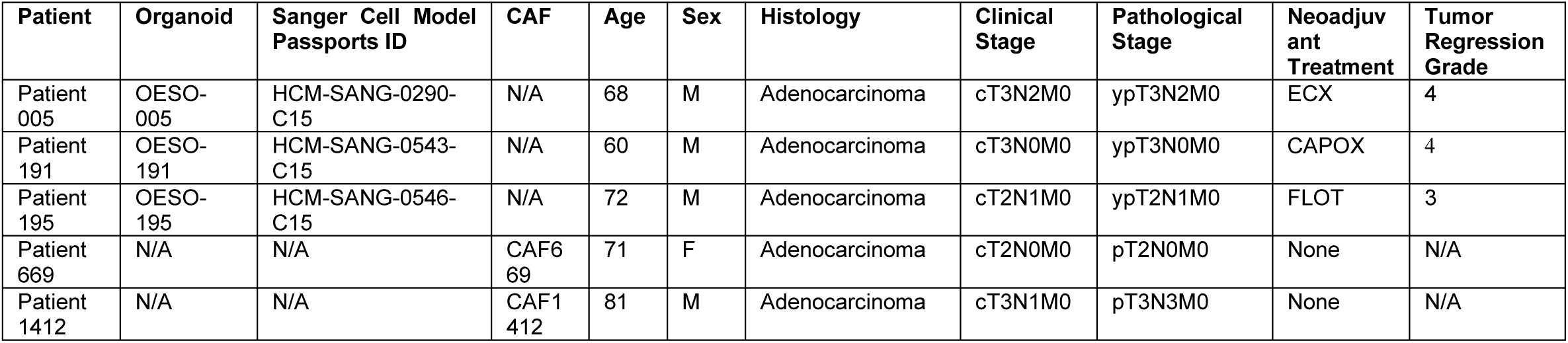
Clinical information for each patient from which primary fibroblasts or organoids were derived and references to associated cell culture and genomic information. ECX – epirubicin, cisplatin and capecitabine; CAPOX – capecitabine and oxaliplatin; FLOT – 5-fluorouracil, leucovorin, oxaliplatin and docetaxel.

### Derivation of Patient-Derived Organoids from EAC Tissue

EAC tumor tissue was removed from the resected specimen during esophagectomy using a sterile 6mm biopsy punch. Tumor tissue was placed in a 50mL falcon tube containing 10mL Hanks’ Balanced Salt Solution (ThermoFisher) on ice for transport. Tissue was taken into a tissue culture hood and washed three times with PBS containing 250ng/mL amphotericin B (Invitrogen) (PBSA) for 10 minutes on ice. Tissue was minced into 1-2mm diameter pieces in digestion buffer with a sterile disposable scalpel: digestion buffer contains EAC-specific organoid media, 30U/mL Collagenase P, 100µg/mL Primocin, 100U/mL penicillin and 100µg/mL streptomycin. Digestion mix was transferred into a 50mL falcon tube and incubated at 37°C with continuous rotation for 1-2 hours until tissue was digested and a cloudy suspension was obtained. Every 15 minutes during the incubation period, the suspension was triturated several times using descending sizes of pipette tip to mechanically break apart the tissue: first 10mL serological pipettes, then 5mL serological pipettes, then 1000µL pipette tips for the last two incubations. The suspension was passed through a 100µm cell strainer and the strainer was washed to collect remaining cells, then centrifuged at 800g for 2 minutes to pellet. Suspension was washed twice with 30mL PBS and centrifuged to dilute out digestion enzymes. The cell pellet was gently resuspended in 200µL of BME2, deposited as droplets on a single well of a 6-well plate, and allowed to set at 37°C for 15 minutes. EAC-specific organoid media consisted of advanced DMEM/F12 containing 10mM HEPES, GlutaMAX, N-2 and B27 supplements, 100ng/ml noggin, 1.25mM N-acetyl-cysteine, 10mM nicotinamide, 50ng/mL EGF, 500nM A83-01, 3µM SB202190 and 100ng/mL FGF-10 as previously described^15^. Conditioned media was collected from L-Wnt3a cells (ATCC CRL-2647™) and Cultrex HA-R-Spondin1-Fc 293T cells (R&D Systems), and the final organoid media formulation contained 20% (v/v) R-Spondin1 conditioned media and 50% (v/v) L-Wnt3a conditioned media. 100U/mL penicillin, 100µg/mL streptomycin and 250ng/mL amphotericin B were added for initial organoid expansion. Once the organoids become crowded or dark, they were passaged by detaching with a pipette, washing with PBS and centrifuging at 800x g for 2 minutes. The pellet is resuspended in TrypLE for 10 minutes at 37C and then centrifuged, resuspending in gel containing 320µL BME2 and 80µL cold EAC organoid growth media and deposited as droplets in two wells of a 6-well plate as before. From this point onwards, antibiotic and antimycotic were withdrawn from the organoid growth media.

### Derivation of Primary Fibroblasts from EAC Tissue

As above, EAC tumor tissue was removed from the resected specimen during esophagectomy using a sterile 6mm biopsy punch. Tumor tissue was placed in a 50mL falcon tube containing 10mL Hanks’ Balanced Salt Solution (ThermoFisher) on ice for transport. Tissue was taken into a tissue culture hood and washed with PBS containing 250ng/mL amphotericin B (PBSA) for 10 minutes on ice. Tissue was minced into 1-2mm diameter pieces in a fresh change of PBSA with a sterile disposable scalpel. Using a scalpel, wells of 6-well plates were scratched in an ‘X’ pattern to create grooves in the plastic, and a single piece of tissue per well was tucked into the middle of the grooves to secure it to the plastic. DMEM containing 10% fetal calf serum, 4mM L-glutamine, 100U/mL penicillin, 100µg/mL streptomycin and 250ng/mL amphotericin B was gently overlayed and media was changed twice weekly until fibroblasts grew out of the tissue and onto the surrounding plastic. Once this occurred, the tissue piece was gently detached using a 200µl pipette tip. Fibroblasts were gently washed using sterile PBS, detached using TrypLE (ThermoFisher), pooled into a single T25 tissue culture flask, and passaged into larger flasks when 80% confluent. At this point, primary fibroblasts were passaged at a ratio of 1:2-1:3 depending on growth rate and were grown without the addition of amphotericin B to the media.

### Production of Assembloid Models

Assembloid methodology was adapted from pancreatic cancer co-culture methodology previously published by Seino et al^23^. To set up assembloids, a single well of a 6-well plate containing organoids and two T175 flasks of 80% confluent fibroblasts are required. In our experience this takes 2-3 weeks depending on the growth rate of the fibroblasts. Fibroblasts were detached from the flask by washing in PBS and incubating with TrypLE for 10 minutes, tapping the flask at the end of incubation to facilitate detachment. Fibroblasts were collected by washing in complete DMEM, centrifuging at 800g for 5 minutes. The fibroblast pellet was resuspended in 5mL of PBS and counted using a hemocytometer. Meanwhile, organoids were collected by pipetting the domes off the well plastic with pre-existing media, washing the well with PBS to collect remaining organoids, pooled and centrifuged at 800g for 2 minutes. Organoids were resuspended in 5mL of TrypLE and incubated at 37°C for 10-15 minutes, until organoid structures disaggregated into single cells and small cell clusters. Organoids were centrifuged as before and resuspended in 5mL PBS for counting.

For each model to be made, 2.5×10^5^ organoid cells and 5×10^5^ fibroblasts were mixed together into a single falcon tube. 10% BME2 in ice cold complete DMEM was prepared, allowing 100µL per model. Cell suspension was centrifuged at 800g for 2 minutes and the cell pellet was resuspended in 10% BME2. 100µL of cell suspension was dispensed into each well of a Nunclon Sphera low attachment 96-well plate. The plate was centrifuged at 400g for 3 minutes at room temperature to facilitate cell aggregation, then incubated at 37°C overnight.

The following day, an extracellular matrix (ECM) gel was prepared containing: 10% filter-sterilized 10X DMEM salt solution, 10% fetal calf serum, 60% rat tail collagen I, 20% BME2. 20µL of ECM gel was prepared per model. Using a sterile disposable 3mL Pasteur pipette, co-cultures were picked from each well of the 96-well plate and deposited in a single drop onto tissue culture plastic while minimizing liquid carryover, either into Ibidi 8-well chamber slides or 96-well plates. Excess media around the co-culture was carefully aspirated using a 20µL pipette, and then a single 20µL drop of ECM gel was overlayed onto the co-culture. The tissue culture vessel was inverted to discourage co-culture attachment to the bottom surface and placed in a 37°C incubator for 15 minutes to allow the ECM gel to set. Once set, 200µL of prewarmed complete DMEM was gently overlayed. Co-cultures were grown this way for another 7 days, with growth media being changed completely on day 4 of culture.

### Wholemount Immunofluorescence on Assembloid Models

Wholemount immunofluorescence methodology was adapted from Dekkers et al^31^ with antigen retrieval methods from the FLASH protocol^32^ added to improve antibody penetration and clearing.

Co-cultures were fixed for 45 minutes in 4% paraformaldehyde in PBS at 4°C, then washed twice in cold PBS. Excess ECM gel was removed using 28G needles under a stereomicroscope, an optional step which improves reagent penetration and improves the working distance available for later confocal imaging. Co-cultures were collected into 1.5mL Eppendorf tubes and epitope retrieval was performed using FLASH reagent 2 (250g/L urea, 80g/L Zwittergent in 200mM boric acid buffer, pH 7.0) with gentle rotation for 2 hours at 37°C. Co-cultures were transferred into 24-well plates, pooling by condition. Co-cultures were washed in organoid washing buffer (OWB) (0.2% BSA and 0.1% Triton X-100 in PBS) at room temperature three times for ten minutes, solution was aspirated and primary antibodies (**Table 2**) diluted in OWB were added to each well (400µL of diluted antibody per well). The plate was incubated for 48 hours at 4°C with gentle rocking. Cocultures were then washed briefly three times with OWB, and then washed three more times for: 30 minutes, 1 hour, and 2 hours respectively. OWB was completely aspirated, secondary antibodies (**Table 2**) were diluted in OWB and DAPI was added to 1µg/mL final concentration. Secondary antibody solution was incubated for 48 hours at 4°C with gentle rocking.

**Table 2.** Antibodies and fluorescent probes used for immunofluorescence.

| Antibody/Reagent | Species origin | Dilution | Antigen Retrieval (for IHC) | Manufacturer | Product Code |
| --- | --- | --- | --- | --- | --- |
| Anti-Pan-Cytokeratin (clone AE1/AE3) | Mouse IgG | 1:100 | Citrate pH 6.0 | Agilent Dako | M351529-2 |
| Anti-Vimentin | Rabbit IgG | 1:200 | Citrate pH 6.0 | Cell Signaling Technology | 5741S |
| Anti-Cytokeratin 7 (clone EPR1619Y) | Rabbit IgG | 1:200 | Tris-EDTA pH 8.0 | Abcam | Ab68459 |
| Anti-Cytokeratin 20 (clone Ks20.8) | Mouse IgG | 1:100 | Tris-EDTA pH 8.0 | Invitrogen | MA5-13263 |
| Anti-Ki-67 | Rabbit IgG | 1:200 | N/A | Abcam | Ab15580 |
| Anti-alpha Smooth Muscle Actin (clone 1A4) | Mouse IgG | 1:200 | Citrate pH 6.0 | Agilent Dako | M085101-2 |
| Anti-Periostin | Rabbit IgG | 1:500 | Citrate pH 6.0 | Abcam | Ab14041 |
| DAPI | N/A | 1µg/mL | N/A | Sigma-Aldrich | D9542-5MG |
| Phalloidin-iFluor594 | N/A | 1:1000 | N/A | Abcam | ab176757 |
| AlexaFluor-488 Anti-Mouse IgG | Goat IgG | 1:500 | N/A | Invitrogen | A-11029 |
| AlexaFluor-633 Anti-Mouse IgG | Goat IgG | 1:500 | N/A | Invitrogen | A-21052 |
| AlexaFluor-488 Anti-Rabbit IgG | Goat IgG | 1:500 | N/A | Invitrogen | A-11008 |
| AlexaFluor-568 Anti-Rabbit IgG | Goat IgG | 1:500 | N/A | Invitrogen | A-11036 |

After repeating the washes as with primary antibody incubations, co-cultures were transferred to fructose-glycerol clearing buffer (60% (v/v) glycerol in 2.5M fructose) for clearing at least 30 minutes at room temperature prior to imaging. Individual co-cultures were spotted onto a glass coverslip with a drop of fructose-glycerol clearing buffer and mounted for imaging in a Leica SP8 laser-scanning confocal microscope. Optical sections were acquired until intensity of DAPI nuclear stain tailed off, normally at ∼200-250µm z-depth. Image files were processed in Imaris v9.5.1 (Oxford Instruments) and rendered in blend mode. Orthogonal projections, maximum intensity z-projections and movies were generated using the same software (**Supplementary movies S1-3**).

### Agarose Embedding of Organoids and Assembloids for Paraffin Histology

Organoids in monoculture and co-culture were grown as described above. For monocultures, organoids were collected by pipetting up and down gently to dissociate the BME2 and collected into a 1.5mL microcentrifuge tube. For co-cultures, specimens were washed and fixed *in situ* on the tissue culture plastic, either on µ-Slide 8-well chamber slides (ibidi) or in 96-well plates. In both cases, cultures were gently washed with ice cold PBS, then fixed in 4% paraformaldehyde for 45 minutes at 4°C. Monoculture organoids were allowed to settle naturally at the bottom of the tube during washes before gently aspirating supernatant.

To handle organoids more easily during paraffin embedding, organoids and co-cultures were pre-embedded in 2% (w/v) agarose. PFA-fixed organoids and co-cultures kept in PBS were allowed to settle to the bottom of the tube and equilibrate to 60°C in a hot block, supernatant was removed, and then 500uL agarose solution pre-equilibrated to 60°C was added. Tubes were placed in a hot block set to 60°C to allow specimens to sink for 10 minutes, and then placed on ice for 5 minutes to solidify the agarose. Agarose blocks were removed from the tubes, trimmed with a scalpel, and placed in 70% ethanol for at least 1 hour prior to processing for paraffin embedding using standard histological protocols.

### Tinctorial Stains

#### Hematoxylin & Eosin Staining

FFPE tissue sections of 4µm thickness were deparaffinized, rehydrated, stained with Mayer’s hematoxylin for 5 minutes, and blued in running tap water. Sections were counterstained with Eosin Y for 5 minutes, rinsed briefly in distilled water, then dehydrated, cleared and mounted in resinous mounting medium. For imaging, slides were scanned using an LM dotSlide (Olympus) with a 20x magnification objective. Scanned H&E stains were reviewed digitally by a specialist gastrointestinal pathologist blinded to specimen identity to assess histological differentiation in both patient tumor tissue and in co-culture specimens.

#### Alcian Blue – Periodic Acid Schiff Staining

FFPE tissue sections of 4µm thickness were deparaffinized, rehydrated, and stained in Alcian Blue (1% Alcian Blue (w/v) in 1% (v/v) aqueous acetic acid). Sections were washed in running tap water for 2 minutes, briefly rinsed in distilled water, treated with 0.5% (w/v) aqueous periodic acid for 5 minutes, rinsed again in distilled water, and then stained with Schiff’s reagent for 10 minutes. Sections were counterstained with Mayer’s hematoxylin for 2 minutes, blued in tap water, dehydrated, cleared and mounted in resinous mounting medium. For imaging, slides were scanned using an Olympus LM dotSlide with a 20x magnification objective.

#### Picro-Sirius Red Staining

FFPE tissue sections of 4µm thickness were deparaffinized, rehydrated, stained with Mayer’s hematoxylin for 2 minutes, then stained with 0.1% (w/v) Sirius Red F3B in saturated aqueous picric acid for 1 hour. Sections were rinsed for 30 seconds in 0.01% (v/v) HCl, rinsed in distilled water, dehydrated, cleared and mounted in resinous mounting medium. For imaging, slides were scanned using an Olympus LM dotSlide with a 10x magnification objective and an analyzer and polarizing filter inserted.

### Double immunofluorescence on FFPE embedded tissue and organoids

FFPE tissue sections of 4µm thickness were deparaffinized and rehydrated in a graded ethanol series. Heat-induced epitope retrieval was performed by microwaving on 50% power for 25 minutes in sodium citrate buffer (10mM sodium citrate, 0.05% Tween-20, pH 6.0) or Tris-EDTA buffer (10mM Tris base, 1mM EDTA, 0.05% Tween-20, pH 9.0) depending on the antibodies to be used (see Table 2). Sections were blocked in 2.5% normal horse serum for 30 minutes and then incubated with primary antibodies at pre-determined dilutions (see Table 2) diluted in antibody buffer (5% bovine serum albumin in TBS with 0.05% Tween-20). Sections were then washed three times, and secondary antibodies diluted in antibody buffer containing 1µg/mL DAPI were applied for 1 hour in the dark. Sections were washed three times and mounted with Mowiol mounting medium^40^, cured overnight at room temperature prior to imaging. Representative images of specimens were acquired using a Leica SP8 laser-scanning confocal microscope.

## Supplementary Data

**Supplementary Movie S1.** Animated 3D rendering of assembloids grown with OES195 and CAF669. Related to Figure 3.

**Supplementary Movie S2.** Animated 3D rendering of assembloids grown with OES005 and CAF1412. Related to Figure 3.

**Supplementary Movie S3.**Animated 3D rendering of assembloids grown with OES191 and CAF1412. Related to Figure 3.

## End Matter

### Author Contributions

Conceptualization, B.P.S., Z.S.W. and T.J.U.; Methodology, B.P.S. and D.A.J.; Investigation, B.P.S., L.A.N., C.T., D.A.J., R.B., J.H., R.R., and M.R.Z.; Resources, D.A.J. and T.J.U.; Data Curation, O.J.P., C.T.; Writing – Original Draft, B.P.S., C.T., and Z.S.W.; Writing – Review & Editing, B.P.S., C.T., M.R.Z., Z.S.W., and T.J.U.; Visualization, B.P.S., D.A.J., and M.R.Z.; Supervision, T.J.U. and Z.S.W; Project Administration, B.P.S., Z.S.W., and T.J.U.; Funding Acquisition, T.J.U.

## Acknowledgments

The authors thank University Hospital Southampton patients for their participation in this study, Dr Mathew Garnett and the Cellular Generation and Phenotyping laboratory at Wellcome Trust Sanger Institute for organoid derivation, and the Faculty of Medicine Tissue Bank at the University of Southampton for patient material and organoid biobanking. FFPE tissue blocks were curated by Research Histology, Department of Cell Pathology, University Hospital Southampton. Histology was supported by Histochemistry Research Unit, University of Southampton. Bioimaging, including slide scanning and laser-scanning confocal microscopy, was supported by Biomedical Imaging Unit, University Hospital Southampton. Prof. Tim J. Underwood is supported by a Royal College of Surgeons of England and Cancer Research UK Advanced Clinician Scientist Fellowship (A23924). Schematics in figures were created with BioRender.com.

